# Genome report: Genome sequence of *Phymata mystica* (Evans), an ambush bug

**DOI:** 10.64898/2026.02.27.708606

**Authors:** Ethan Grebler, Tracy Liesenfelt, Paul K. Masonick, Andrew J. Mongue

**Affiliations:** Department of Entomology and Nematology, Institute of Food and Agricultural Sciences, University of Florida, 1881 Natural Area Drive, Steinmetz Hall, Gainesville, FL 32611, USA; Florida Museum of Natural History, McGuire Center, 3215 Hull Road, Gainesville, FL 32611, USA

**Keywords:** Phymatinae, Reduviidae, venomics, venom conservation, hematophagy, Hemiptera, genome assembly

## Abstract

Recent advances in sequencing technology have made the sequencing of non-model organisms significantly more streamlined and feasible. Using these technologies, we begin to address the lack of data on non-model organisms, by sequencing the genome of one such species, *Phymata mystica* (Evans 1931), an ambush bug (Hemiptera: Heteroptera: Reduviidae: Phymatinae) specialized for floral sit-and-wait style predation. Our genome assembly is 710 Mb, in which 99.7% of this sequence is assembled into 14 chromosomal scaffolds. We found that repetitive elements accounted for 58.85% of the sequence. We report 26,760 protein-coding genes in a preliminary annotation of the genome. Using these new resources, we explored both macrosynteny and gene conservation. Starting with chromosome structure, we found that *P. mystica* has a single X chromosome, unlike other well-assembled Reduviids in which the X apparently split into two linkage groups. Exploring this new annotation, we found a number of venom proteins conserved between *P. mystica* and the other venomous Heteroptera with reference genomes, primarily serine proteases, metallopeptidase and heteropteran venom family proteins. These results provide a new framework for the evolution of venom in this group of insects and further demonstrate the ease with which non-model species can be studied using modern genomic methods.

## Introduction

Molecular research on the ambush bugs (Reduviidae: Phymatinae), a group of early-diverging assassin bugs (Reduviidae) adapted for sit-an-wait predation, has been taken on less due to historical challenges involved with the sequencing of non-model organisms’ genomes. Early research into ambush bugs was conducted mainly by European naturalists for taxonomic purposes, focusing only on a few key species (Evans 1931). The prey records of ambush bugs include a variety of insect plant pests and pollinators, making ambush bugs of potential interest for biological control and for evolutionary biology as an informative contrast to hematophagous assassin bugs (Balduf 1940). Representative species of Phymatinae have been found on every continent but Oceania and Antarctica (Masonick et al. 2017). *Phymata mystica* can be found throughout much of the southeastern region of the United States, as far north as North Carolina and as far south as the Florida keys (Masonick and Weirauch 2020).

Like other predatory Hemiptera, ambush bugs feed by piercing their prey with their maxillary stylets and sucking out their innards. To aid in this process, ambush bugs are venomous with their venoms serving a dual purpose: to both paralyze and to liquify prey (Walker et al. 2017). In reduviids, venom is stored in three glands: the anterior main gland, the posterior main gland, and the accessory gland (Andrew Allan Walker et al. 2018). The anterior main gland is responsible for the production of paralyzing neurotoxins, the posterior main gland is responsible for the production of digestive enzymes and the accessory gland serves as a water reservoir hosting some “toxin like peptides” (Andrew A. Walker, Mayhew, et al. 2018; Andrew A. Walker, Robinson, et al. 2018). The venom glands in assassin bugs have evolved from ancestral salivary glands (Andrew A. Walker, Robinson, et al. 2018).

The venomous proteins contained in the assassin bug sialome (proteome of the salivary gland) can be cytotoxic, hemotoxic, or neurotoxic (Walker et al. 2017; Rügen et al. 2021; Zdenek et al. 2024). Cytotoxic proteins are those possessing antimicrobial, cytolytic, and hemostasis impairing qualities, or any qualities that disrupt cellular metabolism. Hemotoxic proteins elicit coagulation cascade activating or inhibiting, fibrinolytic, hemolytic, hypotensive, platelet aggregation activating/inhibiting and vasodilating activity. Neurotoxic proteins elicit pre- and post-synaptic damage or disruption to the neuron (Zhou et al. 2022).

The sialome of predatory reduviids is primarily made up of serine proteases whereas the venome of hematophagous reduviids is primarily made up of triabins, with very little protease content (Zdenek et al. 2024). Assassin bug venoms are anticoagulatory, leading to hematophagous species exapting this trait for its advantages for blood feeding and losing the additional painful and membrane-permeating effects of predatory assassin bug venom (Zdenek et al. 2024). The anticoagulant activity of assassin bug venoms is facilitated by several protein families. Anticoagulant activity in predatory assassin bugs is promoted by serine proteases through fibrinogen destruction, in hematophagous assassin bugs this function is performed by lipocalins through thrombin inhibition (Zdenek et al. 2024). Anticoagulant activity in bedbugs (*Cimex lectularius*), close relatives of assassin bugs, is facilitated through apyrases (Valenzuela et al. 1996). Predatory assassin bug venoms cause pain through the cytolytic action of redulysins (Fischer et al. 2023).

Despite the wealth of knowledge for these species’ venoms, existing research has yet to explore the venoms of ambush bugs, one of the earliest diverging lineages of assassin bug. Characterization of these insects’ venoms will add context to the evolution of venoms in the family Reduviidae. To begin adding this context, we present the genome, gene annotation, and preliminary venom ortholog analysis of *Phymata mystica*. Here, we generate the chromosome-level assembly and annotation of *P. mystica’*s genome, the first member of Phymatinae to be sequenced, and provide insights into the diversity and evolution of assassin bugs. Alongside the presentation of *P. mystica*’s genome and annotation, we analyze venom conservation between it and other predatory reduviids, as well as hematophagous reduviids, and non-reduviid heteropteran species. These early results add to the understanding of the evolution of venoms across feeding behaviors and diversification of assassin bugs, and serve to benefit future systematic and evolutionary studies of the group.

## Materials and Methods

### Collection of *Phymata mystica*

We sourced DNA from adult *P. mystica* collected from flowers, primarily goldenrod (*Solidago spp*.), within the Natural Area Teaching Laboratory at the University of Florida, located in Alachua County, Florida (29°38’00.2”N 82°22’04.3”W). Collection took place in early October, 2024. We snap-froze live individuals by placing them directly into a −80 °C freezer where we stored tissue until use for sequencing.

### DNA extraction and sequencing

#### Homogenization and DNA extraction

We first utilized tissue from a single male, splitting the body tissue into two extractions, one containing the head and thorax, the other containing the abdomen. These body tissues were homogenized with a Dounce homogenizer and extracted DNA from homogenized tissue with an Omniprep DNA extraction kit for high-quality genomic DNA (G-Biosciences), following the standard Omniprep base kit protocol with adjustments outlined in (Mongue, Markee, et al. 2024) and (Liesenfelt et al. 2025). This modified protocol, which has successfully generated chromosome-level assemblies of other insect species, included lysing tissue for 15-16 hrs overnight on an incubator at 56 °C. To increase DNA yield, we added 2 μL of 20 mg/mL Mussel Glycogen (Thermo Scientific) to each sample during the precipitation step. After Mussel Glycogen was added to each sample, we allowed DNA to precipitate for 1 hr in a −20°C freezer. In tandem, these modifications to the standard Omniprep protocol allowed us to maximize our DNA yields.

#### Quality control and sequencing

We used a Qubit 4 Fluorometer with Qubit dsDNA 1X Broad Range Sensitivity Assay Kit and a Nanodrop spectrophotometer to check the quantity and quality. We sent both DNA and frozen tissue from an additional two individuals to the Arizona Genomics Institute (Tucson, AZ, USA) for Pacific Biosciences (PacBio) HiFi sequencing on a Revio and for Illumina Hi-C library preparation and sequencing, respectively. Following sequencing of the male specimen, we subsequently repeated extraction and sequencing for a single female sample for sex chromosome identification. For this analysis we sequenced extracted DNA as 150bp paired-end Illumina reads with SeqMatic Incorporated (Fremont, CA, USA).

### Genome size estimation

To estimate the expected genome size, we used Jellyfish v. 2.2.4 (Marçais and Kingsford 2011), to count 21-mers with the parameter -m 21. The resultant data was analyzed using custom scripts in R Studio v. 4.2.0 (R Core Team 2020) to estimate both genome size and the coverage depth of homozygous regions.

### Genome assembly

#### Contig assembly from raw reads

We used the Hifiasm v. 0.18 program (Cheng et al. 2021) to assemble our primary genome with the -l 3 parameter for more aggressive removal of haplotigs and the -hom-cov set from the results of the k-mer coverage discussed in the previous section. After primary assembly, we further purged remaining haplotigs from our assembly using the purge_dups pipeline (https://github.com/dfguan/purge_dups).We assessed the assembled genome for completeness with the hemipteran ortholog dataset (hemiptera_odb10) in BUSCO v. 5.7.0 (Manni et al. 2021).

#### Hi-C scaffolding to resolve chromosomes

To bring our assembly to the chromosome-level, we scaffolded our PacBio primary assembly with our Hi-C sequencing data. We used the Arima Hi-C pipeline (https://github.com/ArimaGenomics/mapping_pipeline) to align Hi-C reads to our primary assembly, as we have successfully in other insects (Mongue et al.). We first used YaHS v. 1.1 (Zhou et al. 2022) to do automated scaffolding. Next, we generated a JBAT file of the initial scaffolded assembly which we visualized as a contact map using Juicebox v. 2.3.0 (Robinson et al. 2018) and manually curated misassemblies. Again we assessed the curated genome for completeness with BUSCO.

Finally, we re-aligned the raw male reads to the curated assembly, to identify the X chromosome via coverage difference compared to the autosomes, following similar methods for other arthropods (Mongue and Baird 2024). To differentiate the X from the Y (if assembled), we separately aligned the female sequencing data to the genome. We then used SAMtools v. 1.9 (Li et al. 2009) to sort alignments and calculate per-scaffold coverage. In males, the X and Y should have roughly half the coverage of the autosomes; in females the X and autosomes should have equal coverage and any Y-linked sequences should have no coverage.

### Repeat masking

Repeat characterization and masking was done as a two step process. First, we used RepeatModeler v. 2.0 (Flynn et al. 2020) to find de novo repeats; the “-LTRStruct” parameter was used to additionally find long terminal repeats. The results of this were run through RepeatMasker v. 4.0.9 (Smit et al. 2019), which generated a soft-masked assembly and returned a summary of the identified repeats. This analysis was completed for *P. mystica*, as well as the publicly available chromosome level genomes of *Cimex lectularius, Panstrongylus geniculatus, Rhynocoris fuscipes* and *Triatoma dimidiata*, accessed from the National Center for Biotechnology Information (NCBI). The other taxa were taken through this step as part of gene annotation to perform synteny analyses described below.

### Gene annotation

Owing to limited tissue availability, we did not generate RNAseq datasets for this study, opting instead to use the BRAKER3 annotation program (Gabriel et al. 2024 Feb 29) to annotate protein coding genes from the soft-masked genome. As evidence, we used concatenated protein sequences from hemipteran BUSCOs. We conducted our analysis of the BRAKER annotated genes with the hemiptera_odb10 database to assess completeness. We estimated completeness using only the longest transcript for each gene, which we parsed using a custom bash script.

### Phylogeny methods

To determine the phylogenetic relationships between our generated *P. mystica* and other members of Reduviidae, we downloaded existing datasets for other assassin bugs and outgroups, as shown in **Table S4**. To analyze the transcriptomic data, we assembled the transcriptomes using Trinity v. 2.9.0 (Grabherr et al. 2011). To extract single-copy orthologs for use in phylogenetic inference, we directly ran BUSCO (Manni et al. 2021) on the genomes.

After this, we used a BUSCO_phylogenomics pipeline to gather single-copy BUSCOs found in a minimum of 75% of our sample species and used FastTree v. 2.1.11 (Price et al. 2010) to generate individual gene trees for each protein sequence with accordance to the Jones-Taylor-Thornton model of evolution (Jones et al. 1992) with the CAT approximation for different rates at each site (Lartillot and Philippe 2004).

Next, we generated a consensus species tree using Accurate Species Tree EstimatoR’s (ASTER) ASTRAL tool v1.22 (Zhang et al. 2025) with the parameter “-R” for more subsampling and placement options, and choosing *C*.*lectularius* to root as the outgroup as it is a known member of a separate family within the same infraorder (Cimicomorpha). We assessed confidence in our generated tree with the local posterior probability using a quartet-based algorithm (Sayyari and Mirarab 2016).

### Synteny analysis

To further explore the evolution of ambush and assassin bugs, we characterized shared synteny among species with chromosome-level assemblies using the MCScanX toolkit 1.0.0 (https://github.com/wyp1125/MCScanX). In brief, it utilizes NCBI’s Basic Local Alignment Search Tool (BLAST) to conduct pairwise protein sequence comparisons to identify putative 1-to-1 orthologs between species, then uses genomic coordinates of those genes to map orthologous regions between genomes. To visualize the synteny between *P. mystica* and available chromosome-level assemblies, we used the SynVisio: Synteny Browser (https://github.com/kiranbandi/synvisio).

### Ortholog analysis

To determine the presence of venom-protein orthologs, we used Proteinortho v. 6.0 (Lechner et al. 2011), a tool that detects orthologous genes via sequence similarity. We compared the *P. mystica* gene models to the previously identified venoms expressed by two predatory assassin bugs (*Ectomocoris sp*. and *Oncocephalus sp*.) (Zdenek et al. 2024), one hematophagous assassin bug (*T. pallidipennis*) (Zdenek et al. 2024), and one hematophagous heteropteran (*C*.*lectularius*) (**Table S2**). These additional species samples had their repeats modeled and masked in the same manner as our *P. mystica* sample.

The identification of venom proteins were taken from venom profiles described in Zdenek et al. 2024 and Hernández-Vargas et al. 2017. The venom proteins described in these papers were searched against the annotated genome of *P. mystica*. The venom class definitions (cytotoxic, hemotoxic, neurotoxic, digestive, synergist) were taken from a review of the relevant literature and UniProt functional annotation (Valenzuela et al. 1998; Assumpção et al. 2008; Yang et al. 2008; Trevisan-Silva et al. 2010; Choo et al. 2012; Zdenek et al. 2024).

## Results and Discussion

### Initial genome size estimates

We received PacBio HiFi long reads in 3 runs from the same sample and library, the first run yielded 20.52 Gb of read data,the second 17.66 Gb and the third 13.05, for a total of 51.25 Gb (raw read data accessions found in **Table 1**). We found 2 peaks in our k-mer coverage plot at a sequencing depth of 14x, representing the heterozygous/hemizygous and homozygous coverage respectively, and estimated the genome to be 741 Mb.

**Table 1.**
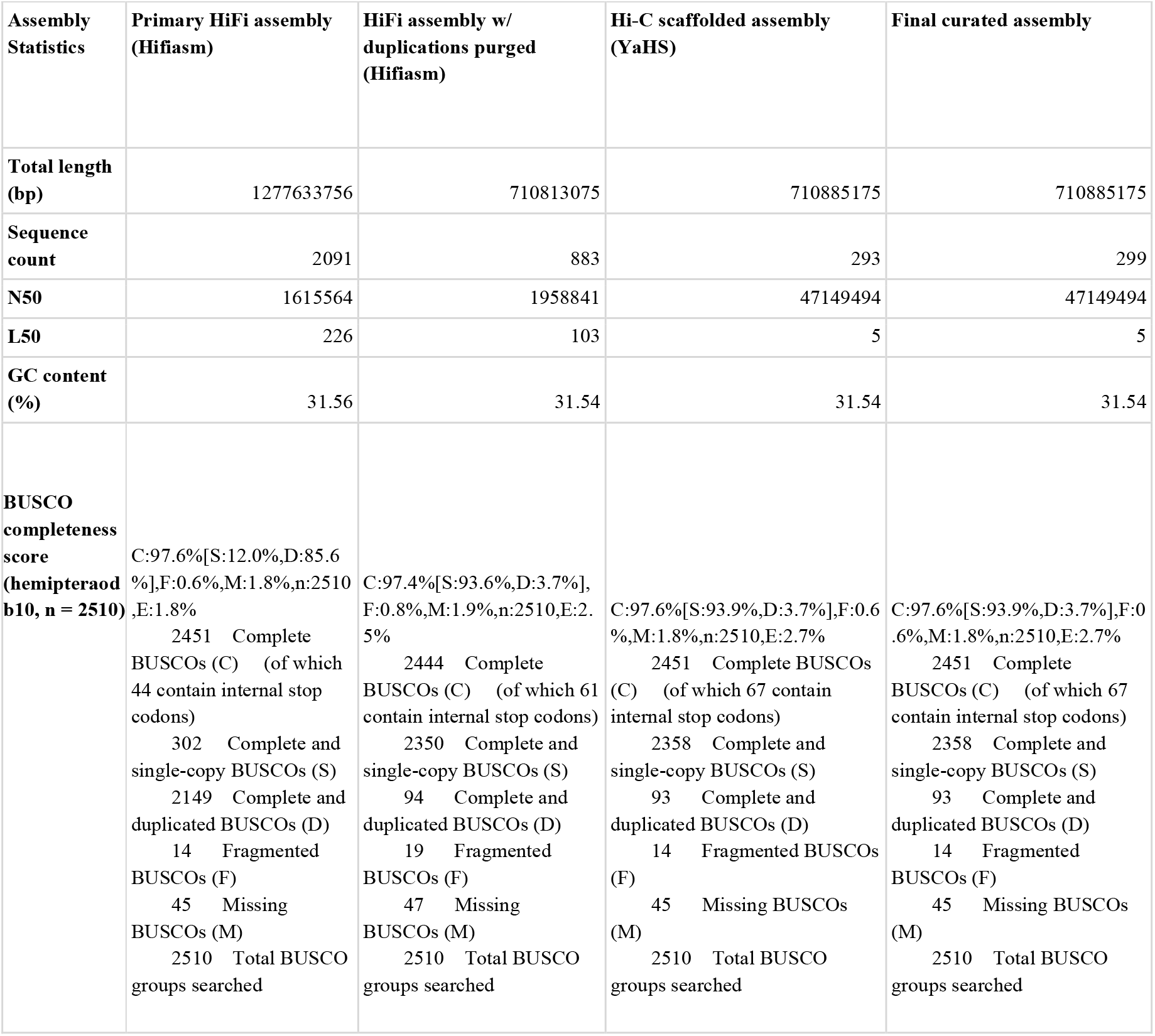
Assembly statistics of the primary HiFi assembly, the HiFi assembly with duplications purged, the Hi-C scaffolded assembly, and the final curated assembly.

| Assembly Statistics | Primary HiFi assembly (Hifiasm) | HiFi assembly w/ duplications purged (Hifiasm) | Hi-C scaffolded assembly (YaHS) | Final curated assembly |
| --- | --- | --- | --- | --- |
| Total length (bp) | 1277633756 | 710813075 | 710885175 | 710885175 |
| Sequence count | 2091 | 883 | 293 | 299 |
| N50 | 1615564 | 1958841 | 47149494 | 47149494 |
| L50 | 226 | 103 | 5 | 5 |
| GC content (%) | 31.56 | 31.54 | 31.54 | 31.54 |
| BUSCO completeness score (hemipteraod b10, n = 2510) | C:97.6% [S:12.0%,D:85.6%],F:0.6%,M:1.8%,n:2510<br>E:1.8%<br>2451 Complete BUSCOs (C) (of which 44 contain internal stop codons)<br>302 Complete and single-copy BUSCOs (S)<br>2149 Complete and duplicated BUSCOs (D)<br>14 Fragmented BUSCOs (F)<br>45 Missing BUSCOs (M)<br>2510 Total BUSCO groups searched | C:97.4% [S:93.6%,D:3.7%],F:0.8%,M:1.9%,n:2510,E:2.5%<br>2444 Complete BUSCOs (C) (of which 61 contain internal stop codons)<br>2350 Complete and single-copy BUSCOs (S)<br>94 Complete and duplicated BUSCOs (D)<br>19 Fragmented BUSCOs (F)<br>47 Missing BUSCOs (M)<br>2510 Total BUSCO groups searched | C:97.6% [S:93.9%,D:3.7%],F:0.6%,M:1.8%,n:2510,E:2.7%<br>2451 Complete BUSCOs (C) (of which 67 contain internal stop codons)<br>2358 Complete and single-copy BUSCOs (S)<br>93 Complete and duplicated BUSCOs (D)<br>14 Fragmented BUSCOs (F)<br>45 Missing BUSCOs (M)<br>2510 Total BUSCO groups searched | C:97.6% [S:93.9%,D:3.7%],F:0.6%,M:1.8%,n:2510,E:2.7%<br>2451 Complete BUSCOs (C) (of which 67 contain internal stop codons)<br>2358 Complete and single-copy BUSCOs (S)<br>93 Complete and duplicated BUSCOs (D)<br>14 Fragmented BUSCOs (F)<br>45 Missing BUSCOs (M)<br>2510 Total BUSCO groups searched |

### Sequencing and assembly

Hi-C scaffolding confidently assembled n=13+X chromosomes, with scaffold_2 identified as the single X via coverage difference from the autosomes in males. The expected Y chromosome may be poorly assembled; we identified nine scaffolds with no coverage in the female sample and non-zero coverage in the male. Of these, only one (scaffold_200) has the expected ∼½ autosomal coverage in males; the rest have very low coverage (**Table S1**) and are likely junk sequences (e.g. error-containing sequences or repeats that differ between samples). The best-supported Y-linked sequence is a mere 24,421bp with nine annotated protein-coding genes. Expanding to include all apparently male-specific sequences yields 199,051bp and a total of 46 genes. By either estimate, the Y is clearly highly degenerated compared to the X, which is the second largest chromosome at 84 Mb and contains 3,478 genes.

More generally, the inferred overall chromosome number is typical of other chromosome-level assemblies of assassin bug genomes (n=12-15), with a key difference being that other sequenced assassin bugs have two distinct X chromosome pairs: X_1_ + X_2_ (Canizales-Silva et al. 2025; Kuznetsova et al. 2025). We placed 99.7% of the assembly within our 14 chromosomal scaffolds. In our final curation steps we made 23 manual adjustments to our assembly using the Juicebox v.2.3.0. tool, including purging 6 duplications and correcting 17 missassemblies. Only adjustments deemed necessary to realign contigs into their correct chromosomal arrangement were made, this was determined through interaction frequency between regions in our assembly. Our final curated assembly received a BUSCO score of 97.6% completeness (93.9% single copy, 3.7% duplicated, 0.6% fragmented and 1.8% missing) (**Table 1**).

Our estimated genome size of 741 Mb was greater than our measured genome size of 710 Mb. Despite having fewer chromosomal scaffolds, the size of *P. mystica*’s genome (720 Mb) is larger than *R. fuscipes* (n=15, 620 Mb) (GCA_040020575.1). Though perhaps not surprising, as sometimes lower chromosome count is associated with larger genome size, e.g. the cottony cushion scale, *Icerya purchasi*, has a genome size > 1 Gb contained in two massive chromosomes (Mongue, Ross, et al. 2024).

### Reduviidae phylogeny

Using available genomic and transcriptomic resources, we explored the phylogenetic relationships between *P. mystica* and other reduviid species, representing 6 subfamilies (**Figure 1**). The tree was rooted using *C. lectularius*, a non-reduviid Hemipteran. We inferred the overall species trees through the use of the hemipteran ortholog dataset (hemiptera_odb10) to build individual gene trees. We found that *P. mystica* is the outgroup to the other sequenced assassin bugs. Our recovered placement of *Panstrongylus geniculatus* and *Hospesneotomae protracta* rendered *Triatoma* paraphyletic. This molecular evidence suggests that the reclassification of these species may be required.

**Fig 1.**
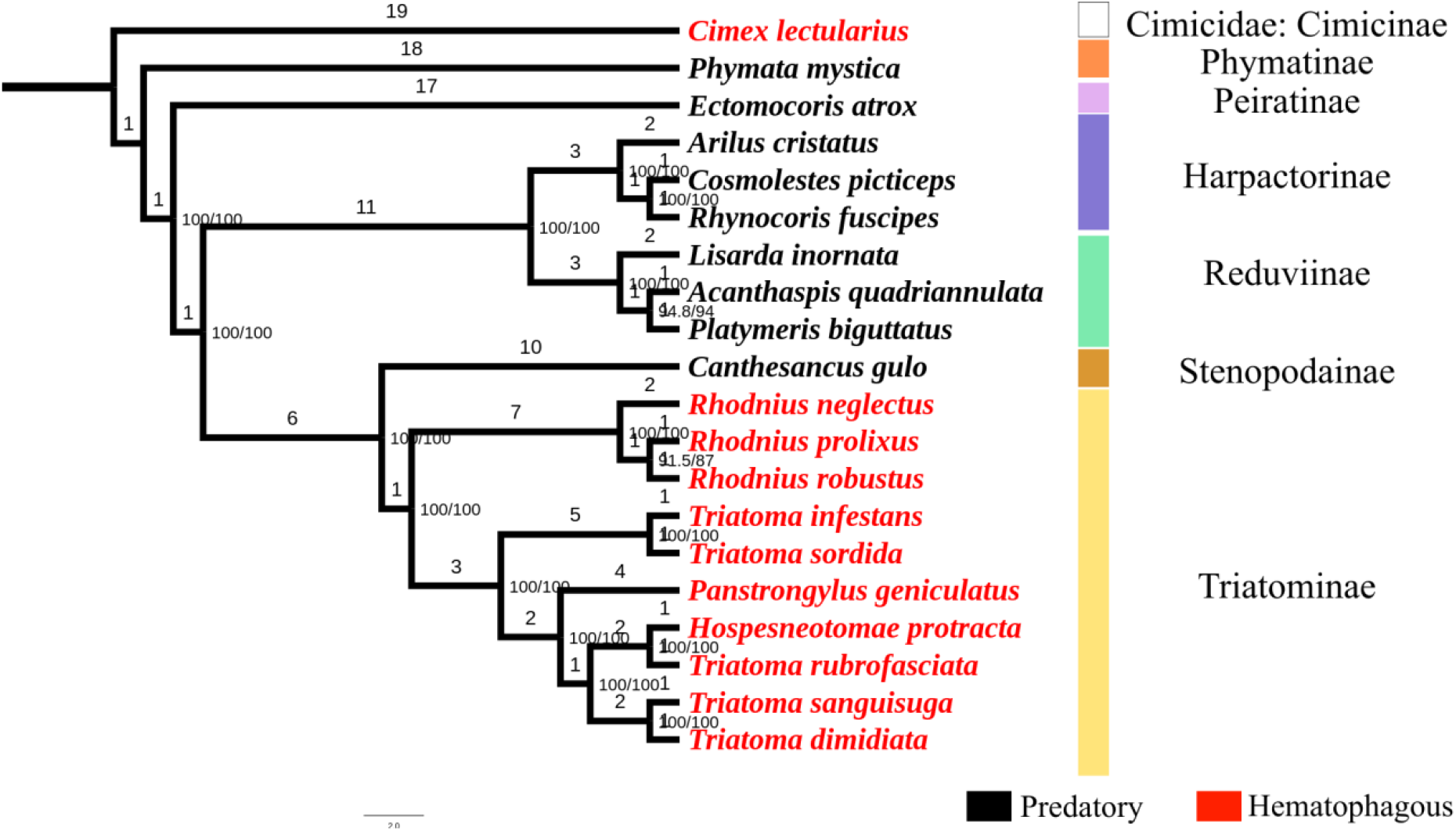
Cladogram of maximum likelihoods for species reported in this study. Support values are formatted as Shimodaira-Hasegawa-like approximate likelihood ratio test (SH-aLRT) / Ultrafast Bootstrap values and are shown at each node. Subfamily designations are listed to the right of their associated terminal(s). Hematophagous species terminals have their names in red.

### Repeat masking

Within our annotated assembly, repeat elements accounted for 58.85% of the sequence (418,374,926 bases). The full extent of repeat content can be seen in **Table 2**. *P. mystica* had the highest repeat content of any reduviid assembled to the chromosome-level, with other species having a repeat element content between 46.26% to 52.55%.

**Table 2.** Summary of the identified and masked repeats in the *P. mystica* genome.

| Category | Count | Length (bp) | Percent |
| --- | --- | --- | --- |
| Retroelements | 516111 | 117,707,204 | 16.56% |
| SINEs | 5962 | 704,433 | 0.10% |
| Penelope | 0 | 0 | 0.00% |
| LINEs | 278,585 | 73,216,923 | 10.30% |
| CRE/SLACS | 0 | 0 | 0.00% |
| L2/CR1/Rex | 92,712 | 25,964,701 | 3.65% |
| R1/LOA/Jockey | 30,801 | 7,748,315 | 1.09% |
| R2/R4/NeSL | 19,188 | 5,453,623 | 0.77% |
| RTE/Bov-B | 33,061 | 9,293,259 | 1.31% |
| L1/CIN4 | 5,233 | 840,251 | 0.12% |
| LTR elements | 231,564 | 43,785,848 | 6.16% |
| BEL/Pao | 3,115 | 2,717,414 | 0.38% |
| Ty1/Copia | 1,303 | 1,038,681 | 0.15% |
| Gypsy/DIRS1 | 136,580 | 25,783,133 | 3.63% |
| Retroviral | 90,400 | 14,220,918 | 2.00% |
| DNA transposons | 335,595 | 74,823,119 | 10.53% |
| hobo-Activator | 89,389 | 19,901,044 | 2.80% |
| Tc1-IS630-Pogo | 108,908 | 23,570,544 | 3.32% |
| En-Spm | 0 | 0 | 0.00% |
| MULE-MuDR | 3,179 | 816,183 | 0.11% |
| PiggyBac | 3,467 | 1,086,840 | 0.15% |
| Tourist/Harbinger | 1,943 | 585,327 | 0.08% |
| Other (Mirage, P-element, Transib) | 694 | 296,324 | 0.04% |
| Rolling-circles | 282,023 | 67,169,590 | 9.45% |
| Unclassified | 1,208,681 | 225,844,603 | 31.77% |
| Total interspersed repeats | — | 418,374,926 | 58.85% |
| Small RNA | 7,788 | 1,725,264 | 0.24% |
| Satellites | 22,492 | 2,632,532 | 0.37% |
| Simple repeats | 263,870 | 14,083,728 | 1.98% |
| Low complexity | 32,170 | 1,719,575 | 0.24% |

### Gene annotation

In total,we identified 26,760 protein-coding genes within our genome for *P. mystica* using the BRAKER3 tool. This annotation showed lower completeness than the genome assembly itself. Once parsed for the longest transcripts of each gene, BUSCO analysis scored the annotation with 92.5% completeness (S:88.8% D:3.7%, 2.5% fragmented and 1.0% missing). *P. mystica* had a larger genome compared to the other predatory reduviid with a chromosome-level sequence, *R. fuscipes*, with only 16,189 protein-coding genes. *P. mystica*’s genome shared a similar number of protein-coding genes to the two hematophagous reduviids with chromosome-level assemblies, 21,125 genes in *T. pallidipennis* and 24,047 in *P. geniculatus*. Future generation of an RNAseq dataset will help refine gene predictions for *P. mystica*.

### Synteny

Our synteny plot found multiple fission events, including one involving the X chromosome after the split between *P. mystica* and higher Reduviidae (**Figure 2**). Despite this change, both parts of the formerly single X are sex-linked in

**Fig 2.**
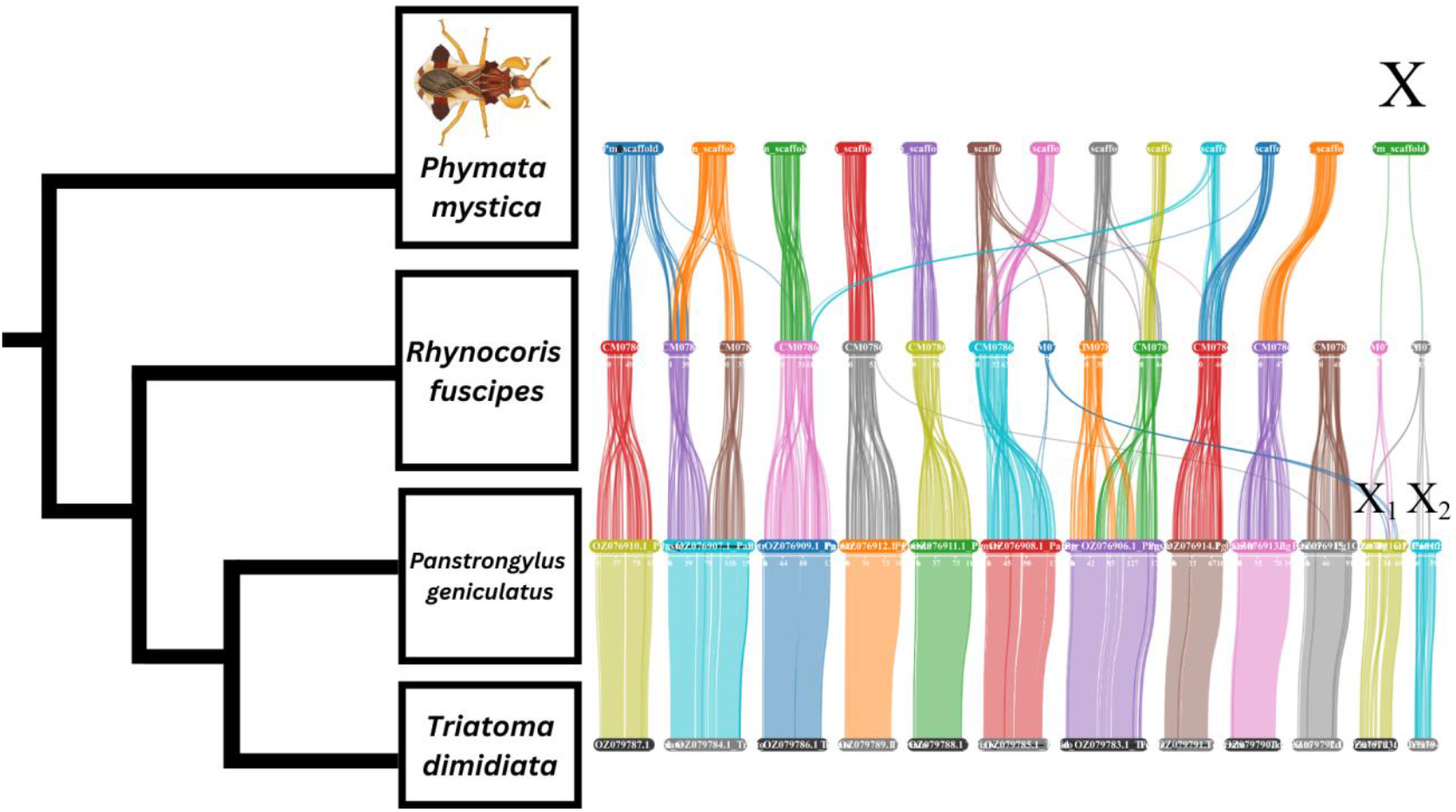
Synteny between chromosome-level assemblies of Reduviidae, showing the fission of the X chromosome and other autosomes after the split between *P. mystica* and the higher Reduviidae. The chromosomes of each of the four species shown are illustrated as colored tabs, lines emanating from these tabs top-down show the movement of genes between chromosomes. The X chromosome is the rightmost tab in *P. mystica*, and the two rightmost tabs as the X chromosome is split into X_1_ and X_2_. The X chromosome is only putatively identified in *P. mystica* and *P. geniculatus. P. mystica* illustrated by TL.

*P. geniculatus*, which suggests that the X_1_X_2_ system is conserved in *Rhynocoris* and *Triatoma*. Several other crossovers, fission and fusion events were found to have occurred in the autosomes in the splits between Phymatinae (*P. mystica*) and Harpactorinae (*R. fuscipes*), and Harpactorinae and Triatominae (*P. geniculatus)*. We found no crossovers, fission or fusion events in the split between *P. geniculatus* and *T. pallidipennis*, both members of Triatominae.

### Ortholog conservation

Our ortholog analysis found 34 unique proteins conserved across the venom proteomes of *C. lectularius, Ectomocoris sp*., *Oncocephalus sp*., *T. pallidipennis*, and the full annotation of *P. mystica*. We found representatives from 19 venom protein families conserved in *P. mystica*. A breakdown of protein families by predicted venomic effect can be seen in **Figure 3**. The largest group represented in our analysis are proteases, mainly serine proteases with some metallopeptidases. Of the serine proteases detected, 8 contained a CUB domain. There is a considerable amount of orthology with proteins in ungrouped heteropteran venom families, including families 1, 2, 3 and 6. The presence of serine proteases and heteropteran venom family proteins in the venom of *P. mystica* is consistent with the sialome of other predatory reduviids (Zdenek et al. 2024). We found two orthologs of lipocalin (Zdenek et al. 2024), the primary component of *T. pallidipennis* venom, in *P. mystica* from both *C. lectularius* and *T. pallidipennis*. A comprehensive list of conserved venoms can be found in **Table S3**. In summary, we found all classes of assassin bug venom groups in the ambush bug, demonstrating that these proteins evolved very early in the family, most likely present in the most recent common ancestor to the reduviids. In contrast, we found more lipocalin family proteins in *P. mystica*, than in *Ectomocoris sp*. and *Oncocephalus sp*., showing changes to venom composition in the outgroup to all other Reduviidae (Phymatinae) despite overall similar content.

**Fig 3.**
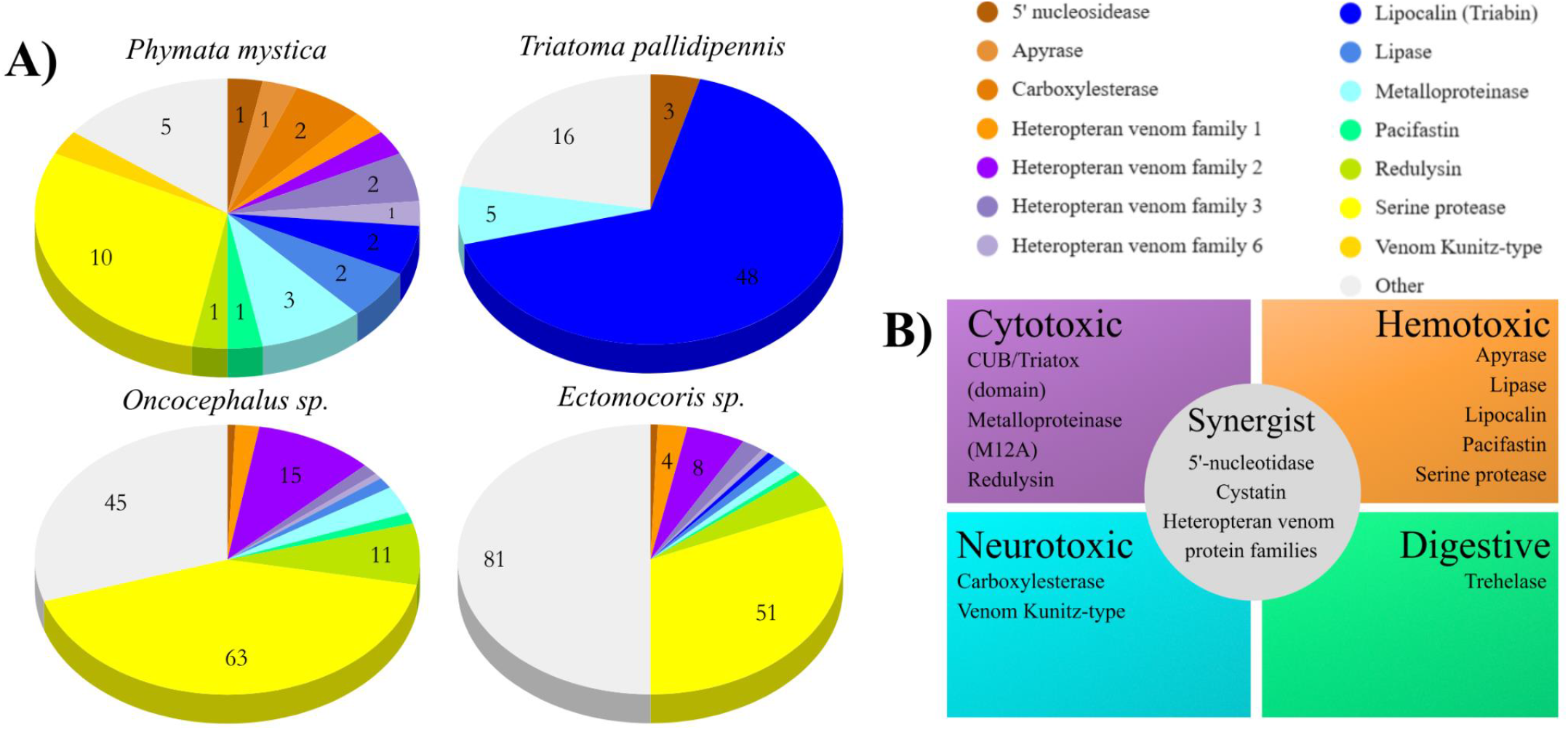
**A)** Venom profile of *P. mystica* inferred from direct orthology, compared with the full venom profiles of related species. The profile inferred for *P. mystica* represents the conserved/ancestral state. Venom class definitions come from a review of relevant literature and uniprot functional annotation as described in the methods section. Lipocalin is present in higher amounts in the inferred *P. mystica* profile than other predatory species. **B)** Protein families inferred to be produced by *P. mystica* from genetic data, grouped by documented venomic effect from arthropod samples. All venom effect classifications are found in *P. mystica*.

## Conclusions

We report a chromosome-level genomic assembly of *Phymata mystica*, a venomous ambush bug in the understudied Phymatinae. This assembly provides the essential resources for further genetic research into this subfamily. Our curated assembly, annotation, and repeat data provide a means of comparison to other members of the family Reduviidae. The record of conservation of venoms produced by this study serves as a jumping off point for more complete venom profiles to be generated from *P. mystica* and further comparisons to elucidate the evolution of venoms across assassin bugs.

## Data Availability

All relevant data will be publicly accessible through the National Library of Medicine: National Center for Biotechnology Information (NCBI). For review purposes pleas use the following link: https://dataview.ncbi.nlm.nih.gov/object/PRJNA1405667?reviewer=4l8jgjfm7lpm4vhdfbj16megk4 Raw sequence data and accession numbers are available in Table 4. The annotation can be found [embargoed until publication], and full venom conservation details are provided as a Supplementary file. Supplemental material available at G3 online.

## Acknowledgements

We acknowledge the UFIT Research Computing for providing computational resources and support that have contributed to the research results reported in this publication. URL: http://www.rc.ufl.edu.

---

**Table S1.**
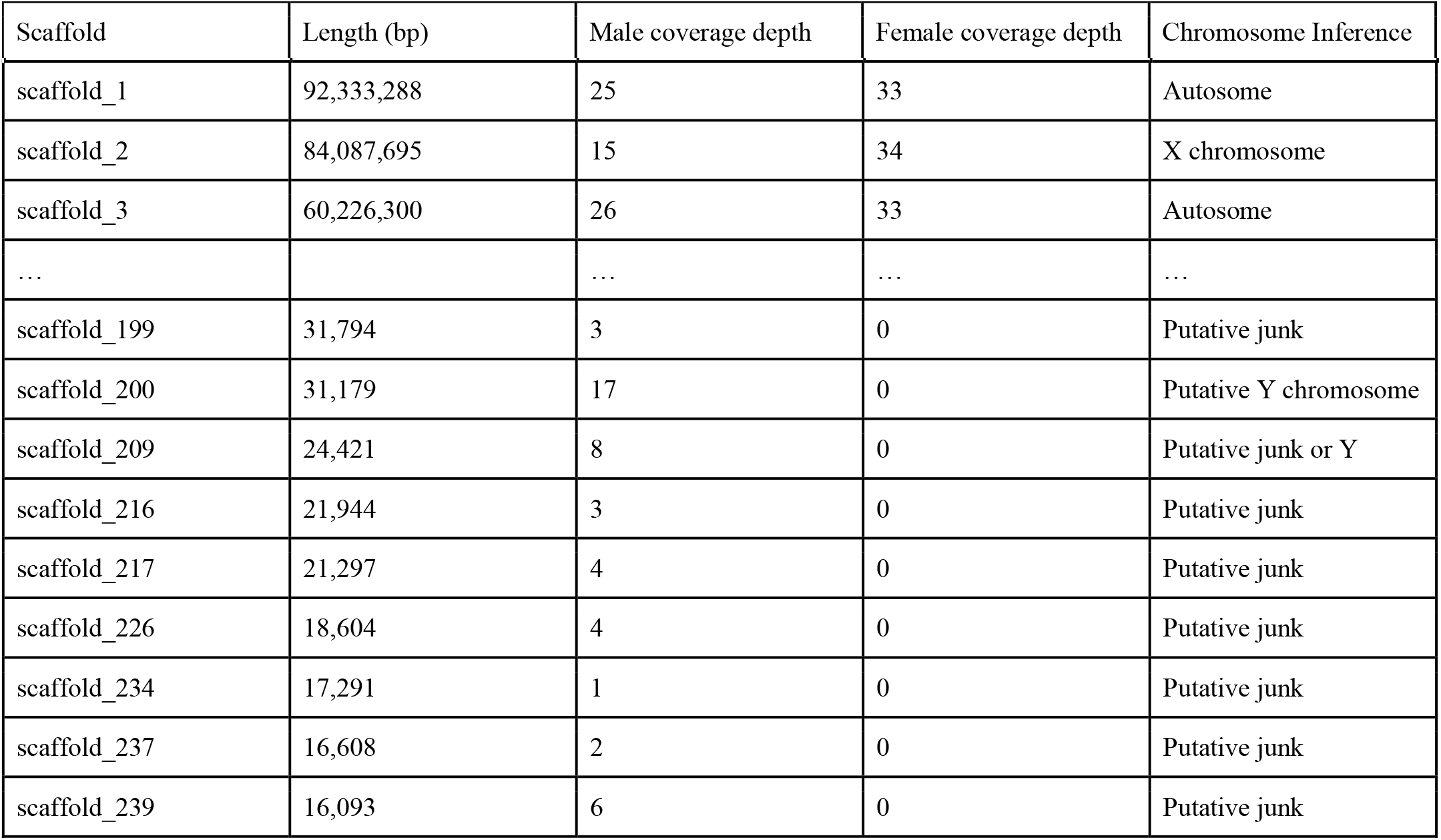
Abridged coverage based inference of sex-linked sequences. For brevity, we show only two autosomal scaffolds to demonstrate the pattern, but all omitted scaffolds had autosomal coverage patterns.

| Scaffold | Length (bp) | Male coverage depth | Female coverage depth | Chromosome Inference |
| --- | --- | --- | --- | --- |
| scaffold_1 | 92,333,288 | 25 | 33 | Autosome |
| scaffold_2 | 84,087,695 | 15 | 34 | X chromosome |
| scaffold_3 | 60,226,300 | 26 | 33 | Autosome |
| ... |  | ... | ... | ... |
| scaffold_199 | 31,794 | 3 | 0 | Putative junk |
| scaffold_200 | 31,179 | 17 | 0 | Putative Y chromosome |
| scaffold_209 | 24,421 | 8 | 0 | Putative junk or Y |
| scaffold_216 | 21,944 | 3 | 0 | Putative junk |
| scaffold_217 | 21,297 | 4 | 0 | Putative junk |
| scaffold_226 | 18,604 | 4 | 0 | Putative junk |
| scaffold_234 | 17,291 | 1 | 0 | Putative junk |
| scaffold_237 | 16,608 | 2 | 0 | Putative junk |
| scaffold_239 | 16,093 | 6 | 0 | Putative junk |

**Table S2.**
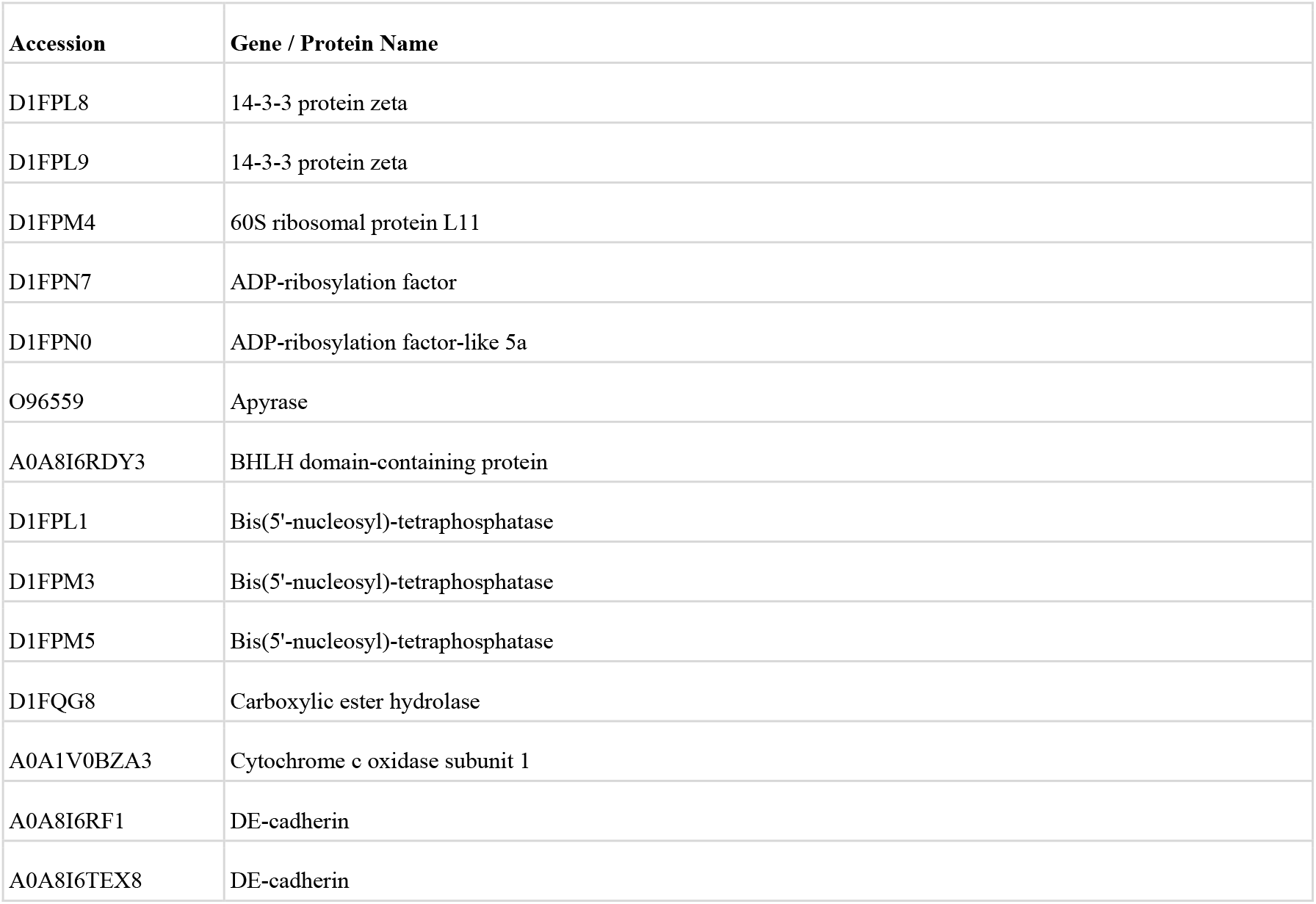
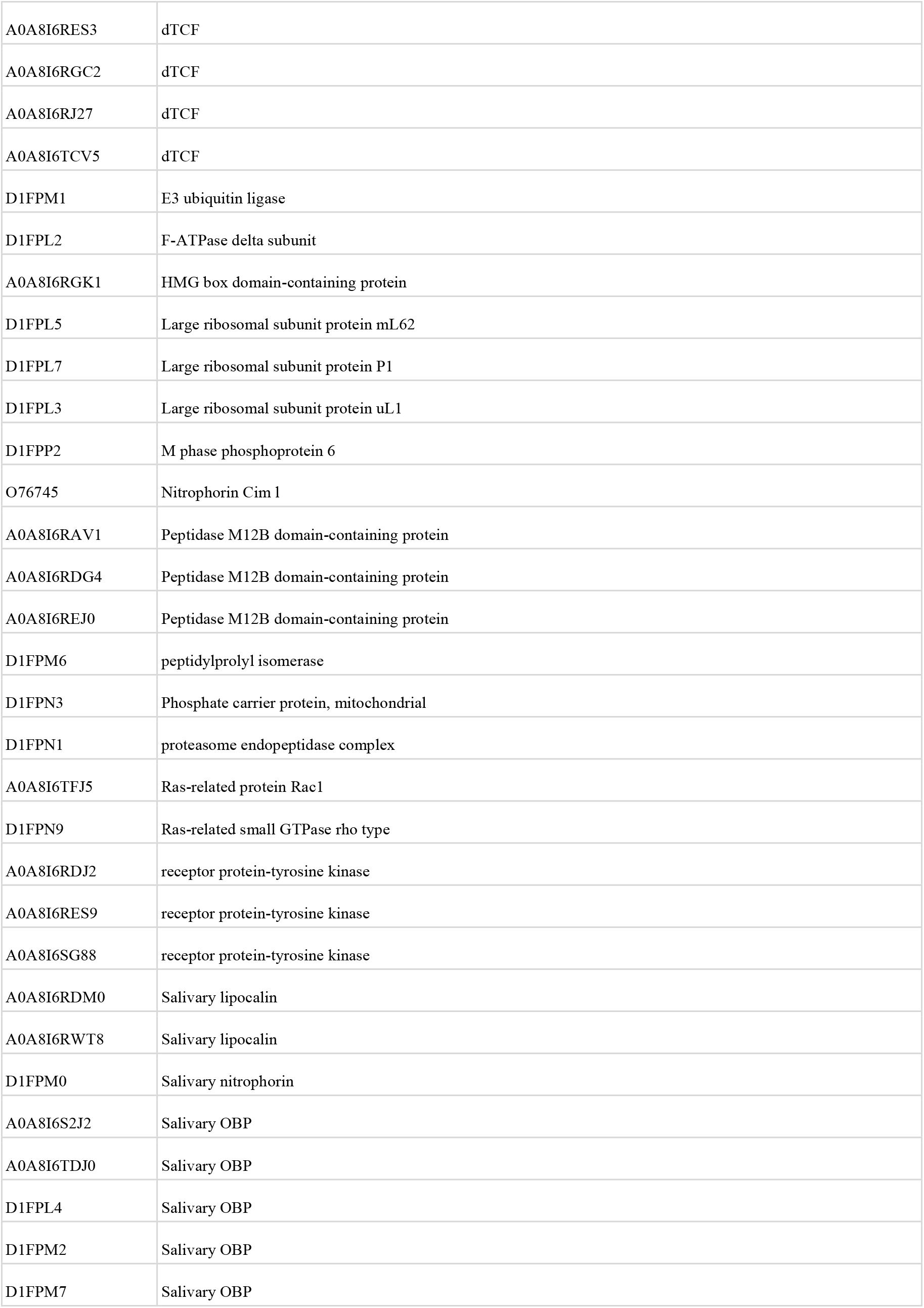
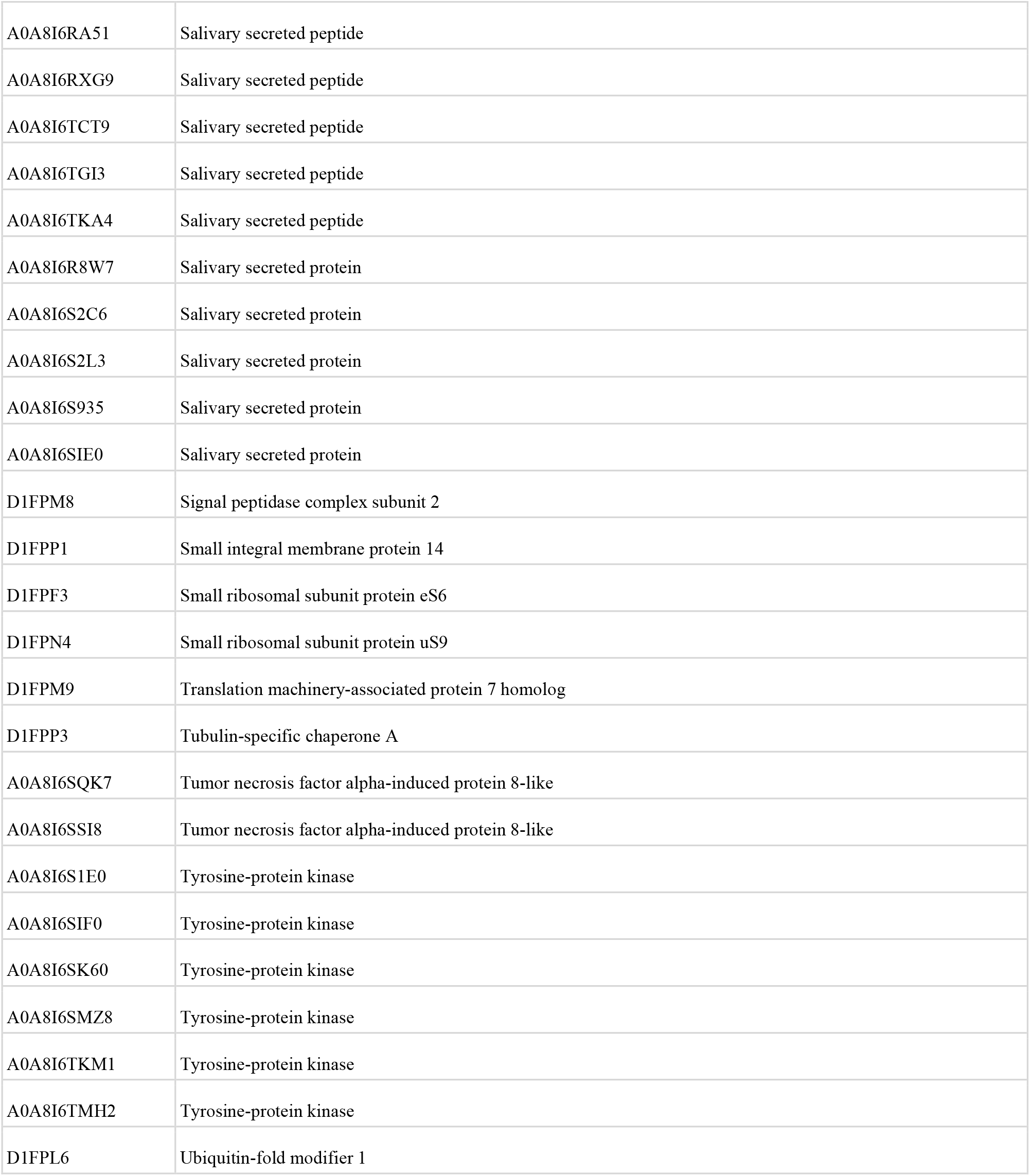
Salivary proteins of *Cimex lectularius* obtained from uniprot based on functional annotation, that were searched against the annotated *Phymata mystica* genome.

**Table S3.**
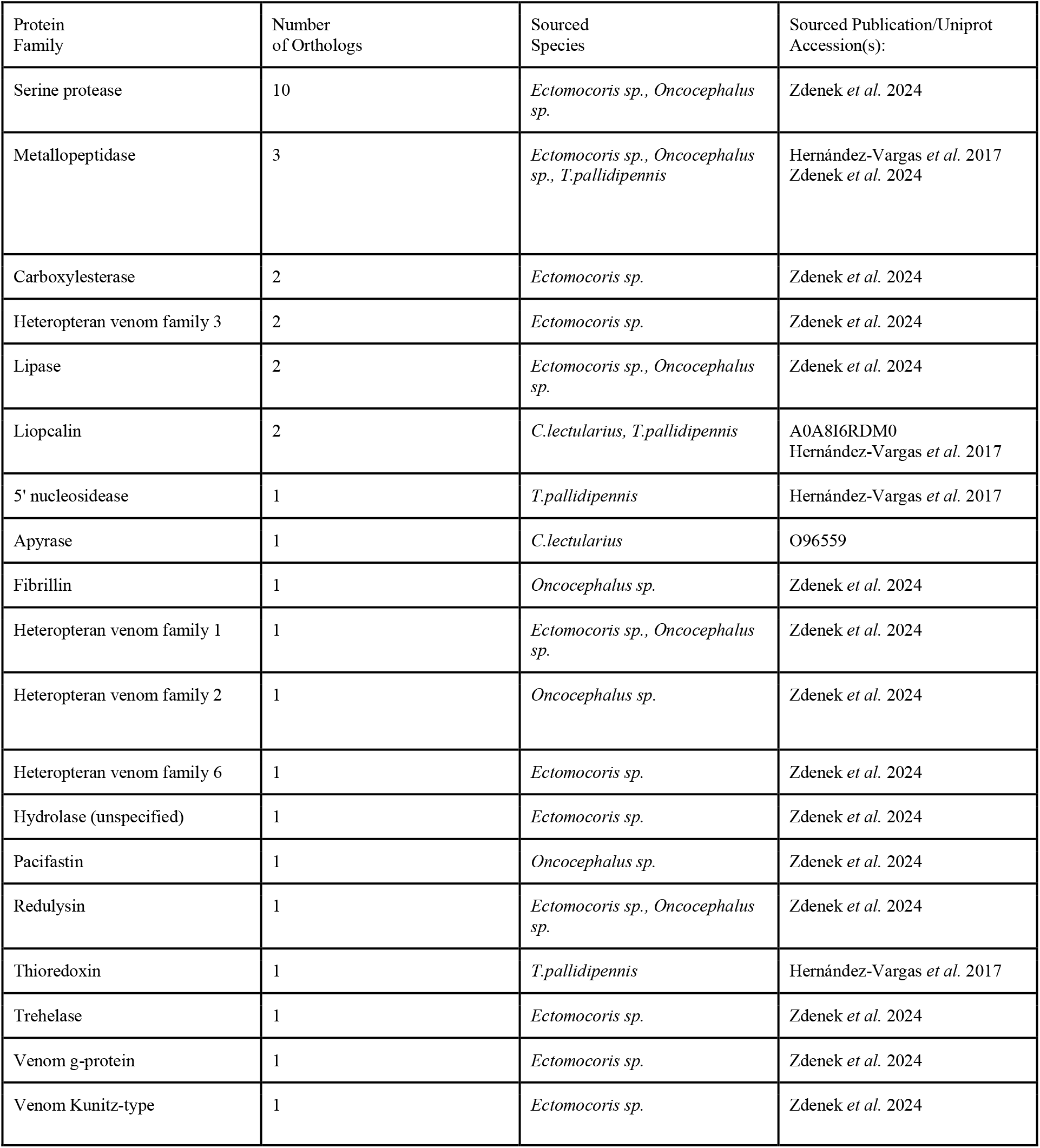
Venom orthologs found in the annotated *Phymata mystica* genome, along with the amount of orthologous proteins found, their sourced organism and sourced publication.

**Table S4.**
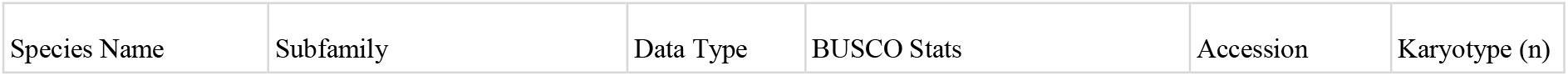
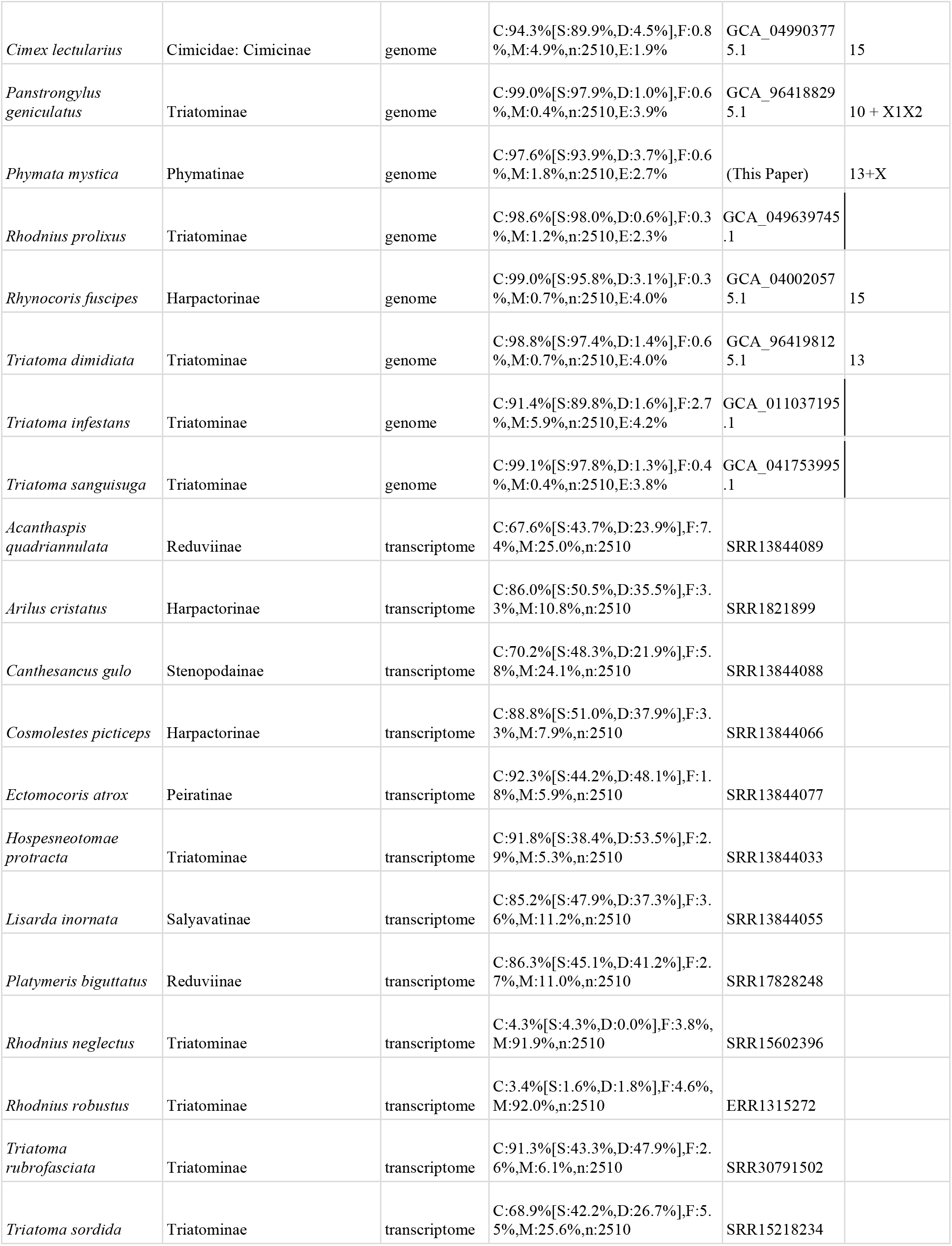
BUSCO Stats for genomic and transcriptomic data used in this study, along with their accession numbers and documented karyotype. If a venom profile was accessed for a species, the publication in which the profile was noted is documented.

